# Energy Landscape of Ubiquitin is Weakly Multidimensional

**DOI:** 10.1101/2021.03.28.437368

**Authors:** Balaka Mondal, D. Thirumalai, Govardhan Reddy

## Abstract

Single molecule pulling experiments report time-dependent changes in the extension (*X*) of a biomolecule as a function of the applied force (*f*). By fitting the data to one-dimensional analytical models of the energy landscape, the hopping rates between the folded and unfolded states in two-state folders, the height and the location of the transition state (TS) can be extracted. Although this approach is remarkably insightful, there are cases for which the energy landscape is multidimensional (catch bonds being the most prominent). To assess if the unfolding energy landscape in small single domain proteins could be one dimensional, we simulated force-induced unfolding of Ubiquitin (Ub) using the coarse-grained Self-Organized Polymer-Side Chain (SOP-SC) model. Brownian dynamics simulations using the SOP-SC model reveal that the Ub energy landscape is weakly multidimensional (WMD) governed predominantly by a single barrier. The unfolding pathway is confined to a narrow reaction pathway that could be described as diffusion in a quasi 1D *X*-dependent free energy profile. However, a granular analysis using the *P_fold_* analysis, which does not assume any form for the reaction coordinate, shows that *X* alone does not account for the height, and more importantly, the location of the TS. The *f*-dependent TS location moves towards the folded state as *f* increases, in accord with the Hammond postulate. Our study shows that, in addition to analyzing the *f*-dependent hopping rates, the transition state ensemble must also be determined without resorting to *X* as a reaction coordinate in order to describe the unfolding energy landscapes of single domain proteins, especially if they are only WMD.

**TOC Graphic:** 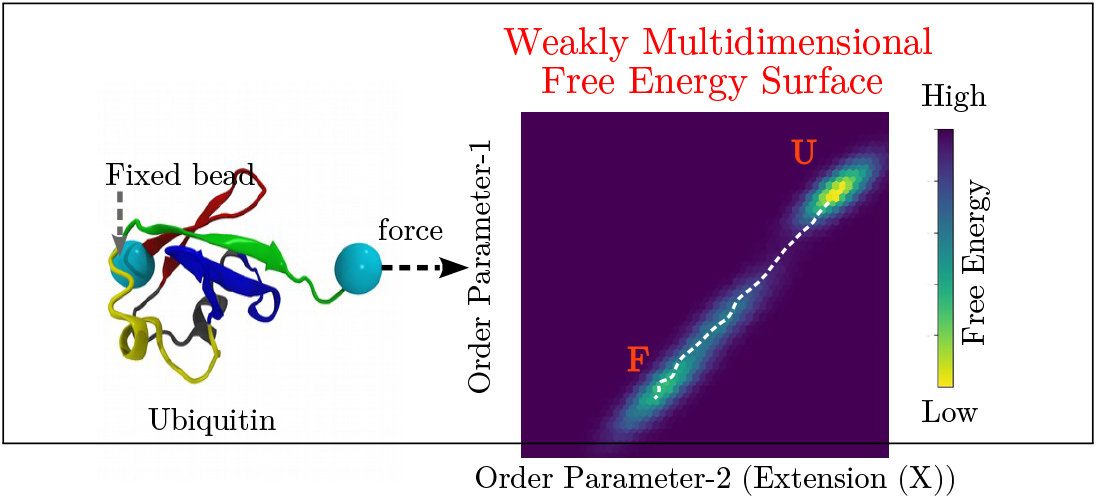

## Introduction

Many cellular functions are triggered by proteins that respond to mechanical stress. ^1,2^ In certain cellular activities, such as muscle contraction and relaxation, cell adhesion, and transport of proteins through a membrane, proteins partially unfold when subject to an external force.^3–5^ These examples illustrate the need to quantitatively understand the molecular basis of the response of biomolecules to forces. Single molecule pulling experiments (optical tweezers, magnetic tweezers, or atomic force microscope (AFM)), in the constant force mode, measure the extension of the molecule as a function of time, which could be used to sort out the structural transitions in a protein. In the force-ramp experiments,^6–11^ force is increased as a function of time by stretching the protein at a constant speed. Typically, but not always, force is applied to the ends of the protein of interest. Pulling at a constant force or speed unfolds the protein, leading to conformational transitions, and evenntually complete unfolding at sufficiently large forces. In the constant force mode (force clamp), structural transitions are detected by sudden changes in molecular extension (*X*) monitored in real time.^12^ By performing the experiments at different values of force *f*, the force-dependent unfolding rate constant *k_u_*(*f*) can be extracted. In the force-ramp experiments,^13^ the protein ends are pulled at a constant speed (or loading rate), and as *f* increases, the extension *X* can be measured. Abrupt changes in the force-extension plots (rips) signify the structural transitions in the protein. In each realization of the experiments, the unfolding force varies stochastically. By performing a large number of experiments, the distribution of rupture forces *p_u_*(*f*) are measured. In principle, the data from the force clamp and constant loading rate modes produce the same information.

The measurements using single molecule pulling experiments have provided a quantitative estimate of the free energy profile (referred to as free energy surface (FES)) by measuring the histogram of *X*. Phenomenological and analytical models based on Kramer’s theory^14^ are routinely used to analyze the data assuming that the folding FES is one-dimensional with *X* as a reasonable reaction coordinate. The Bell model^15^ assumes that the force-dependent unfolding rate *k_u_*(*f*) decreases exponentially with the applied force *f*. The fit of the data to the Bell model may be used to extract the intrinsic unfolding rate *k_u_*(0) at *f* = 0, and the transition state (TS) location, Δ*X*^‡^. A limitation of the Bell model is that it does not relate *k_u_*(*f*) to the barrier height Δ*G*^‡^ or to the shape of the 1D FES. It has become clear that the Bell model is valid only for brittle proteins in which TS does not move with *f*.^16–21^ However, to account for ductile systems (Δ*X*^‡^ changes with *f* ^22^), 1D FES (cubic FES or one with a cusp) for which *k_u_*(*f*) could be analytically calculated have proved to be particularly useful.^7,23^

In an influential article, Dudko, Hummer, and Szabo modeled the FES as linear-cubic and harmonic cusp functions of *X*, and derived analytical expressions^24,25^ for *k_u_*(*f*) by using a modified Kramer’s theory.^14^ The unfolding rate *k_u_*(*f*) connects *k_u_*(0) to the free energy properties, Δ*G*^‡^ and Δ*X*^‡^. These quantities are extracted by fitting the data to the theoretical expression. The theory also assumes that FES is one-dimensional function of *X*, which might not always hold. Indeed, experiments and simulations provide evidence that the FES for some biomolecules could be multidimensional, ^26,27^ involving multiple barriers.^28–30^

If the energy landscape were multidimensional, then it could result in the nonlinearity in log *k_u_*(*f*) versus *f* plot in force clamp experiments^19,31,32^ or in the most probable rupture force *f* * versus the logarithm of loading rate log *r_f_*.^28,33–37^ Is the multidimensionality of the unfolding energy landscape generic? There are only a few single domain proteins for which the presence of multiple unfolding pathways, which requires the use of multidimensionality to explain the experimental data, has been clearly documented.^26,30,38–40^ This suggests that at least strong multidimensionality in single domain proteins may not be the norm but cannot be ruled out unless scrutinized carefully. Ubiquitin (Ub) is an excellent model system to probe the validity of the assumption that the energy landscape is one-dimensional due to the availability of extensive single molecule pulling experimental data.^41,42^ To this end, we performed force-clamp and force-ramp pulling simulations of Ub using the coarse-grained Self-organized Polymer-Side Chain (SOP-SC)^43,44^ model, which allows us to use forces that are similar to those used in the experiments. We observed a downward curvature in the log *k_u_*(*f*) versus *f* plots in force clamp simulations, indicating that the FES is weakly multi-dimensional. ^26^ The nonlinearity observed in both the force clamp and force ramp simulation data due to the movement of the TS towards the folded state accords well with the Hammond postulate. The location of the TS extracted from the experiments does not satisfy the *P_fold_* criterion.^45^ Taken together, our work shows that, in addition to fitting experimentally measured rates to theories that assume that the free energy profile is one dimensional, additional tests that do not rely on an assumed reaction coordinate or experimental data, may be needed to understand the force-induced unfolding dynamics.

## Methods

### SOP-SC Model and Simulation Method

Brownian dynamics simulations at temperatures, *T* = 300 K and 332 K are performed^46^ using the SOP-SC model. ^43,44^ The interactions between the side chains are given by the statistical potential.^47^ The SOP-SC model and the simulation methodology^48,49^ are described in detail in the Supplementary Information (SI). The SOP-SC energy function parameters are given in the SI (Tables S1, S2 and S3).

In the force-clamp simulations, we fixed the *N*-terminal end of Ub (backbone bead of residue-1), and applied a constant force *f* to the *C*-terminal end (backbone bead of residue 76). In the force-ramp simulations, the backbone bead of residue-1 is fixed, and the backbone bead of residue-76 is attached to a spring, with a spring constant, *k_s_* = 35 pN/nm. The other end of the spring is moved at a constant velocity *v* to ramp up the force in order to unfold the protein. The loading rate, *r_f_ = k_s_v*, ranges from 8.75 ×10^4^ pN/s to 8.75 ×10^7^ pN/s. The end-to-end distance, *X*, of Ub is the distance between the backbone beads of residues 1 and 76. The structural overlap function,^50^ χ, is defined as 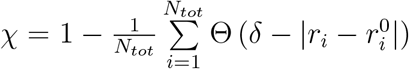, is defined as to distinguish between different populated states of the protein. Here, *N_tot_*(= 11026) number of pairs of interaction centers in the SOP-SC model of Ub assuming that at least two bonds separate the interaction centers, *r_i_* is the distance between the *i^th^* pair of beads, and 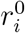 being the corresponding distance in the folded state, Θ is the Heaviside step function, and *δ* = 2 Å.

### Transition state ensemble

The numerically precise transition state ensemble (TSE) is identified from the unfolding trajectories obtained at different *f* values at *T* = 300 and 332 K using the *P_fold_* analysis.^45^ Putative TSE structures are picked from the region in the unfolding trajectories where the folded state starts to unravel. This region satisfies the conditions 0.66 < *χ* < 0.68 and −50.0 kcal/mol < *E_CG_* < −40.0 kcal/mol, where *E_CG_* is the potential energy of a protein conformation in the SOP-SC representation. To compute the commitment probability *P_fold_* for the probable TSEs, 500 short Brownian dynamics simulations are spawned with the friction value *ζ* = 1.0 *m/τ_H_*, where *m* is the mass of the bead, and *τ_H_* is the time unit of the Brownian time step. If the *P_fold_* of a structure is in the range 0.4 < *P_fold_* < 0.6, then the structure is classified as belonging to the TSE.^45^

## Results and Discussion

### Force clamp pulling simulations of Ub

We first performed force-clamp simulations by applying a constant *f* to the protein ends. At *T* = 332 K and *f* = 0 pN, the free energy difference (Δ*G*) between the Ub folded and unfolded states is ≈ 6.6 kcal/mol. The good agreement between the calculated and the experimental values^51^ for Δ*G* provides a measure of validation of the SOP-SC model. Upon application of *f*, the end-to-end distance of the protein, *X* increased by ≈ 22 nm in agreement with experiments,^42^ and previous simulations^52–59^ (Figure S1 and S2).

### Ub unfolding is weakly multidimensional

Force induced unfolding of Ub occurs in a two-state manner without populating any intermediates (Figure S1). At each *f*, we performed 100 independent pulling simulations to compute the average unfolding time ⟨*τ*(*f*)⟩. Assuming two-state kinetics, the force dependent unfolding rate is taken to be *k_u_*(*f*) = ⟨*τ*(*f*)⟩^-1^. The relative location of the TS, Δ*X*^‡^, may be obtained by fitting the simulation data in the [*f*, log(*k_u_*(*f*))] plot to the Bell model, ^15^ 
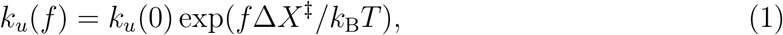
where *k*_B_ is the Boltzmann constant, and Δ*X*^‡^ = *X*^‡^ − *X^F^*, where *X*^‡^ is the end-to-end distance of Ub in the TS, and *X^F^* is the end-to-end distance of Ub in the folded state at *f* = 0 pN. The fit of [*f,* log(*k_u_*(*f*))] to Eq. 1 yields Δ*X*^‡^ ≈ 0.24 ± 0.04 nm, which agrees with the AFM pulling experiment at high forces,^60^ and *k_u_*(0) = 2.1 ± 0.33 s^−1^ (Figure 1). The fitting procedure is described in the SI.

**Figure 1:**
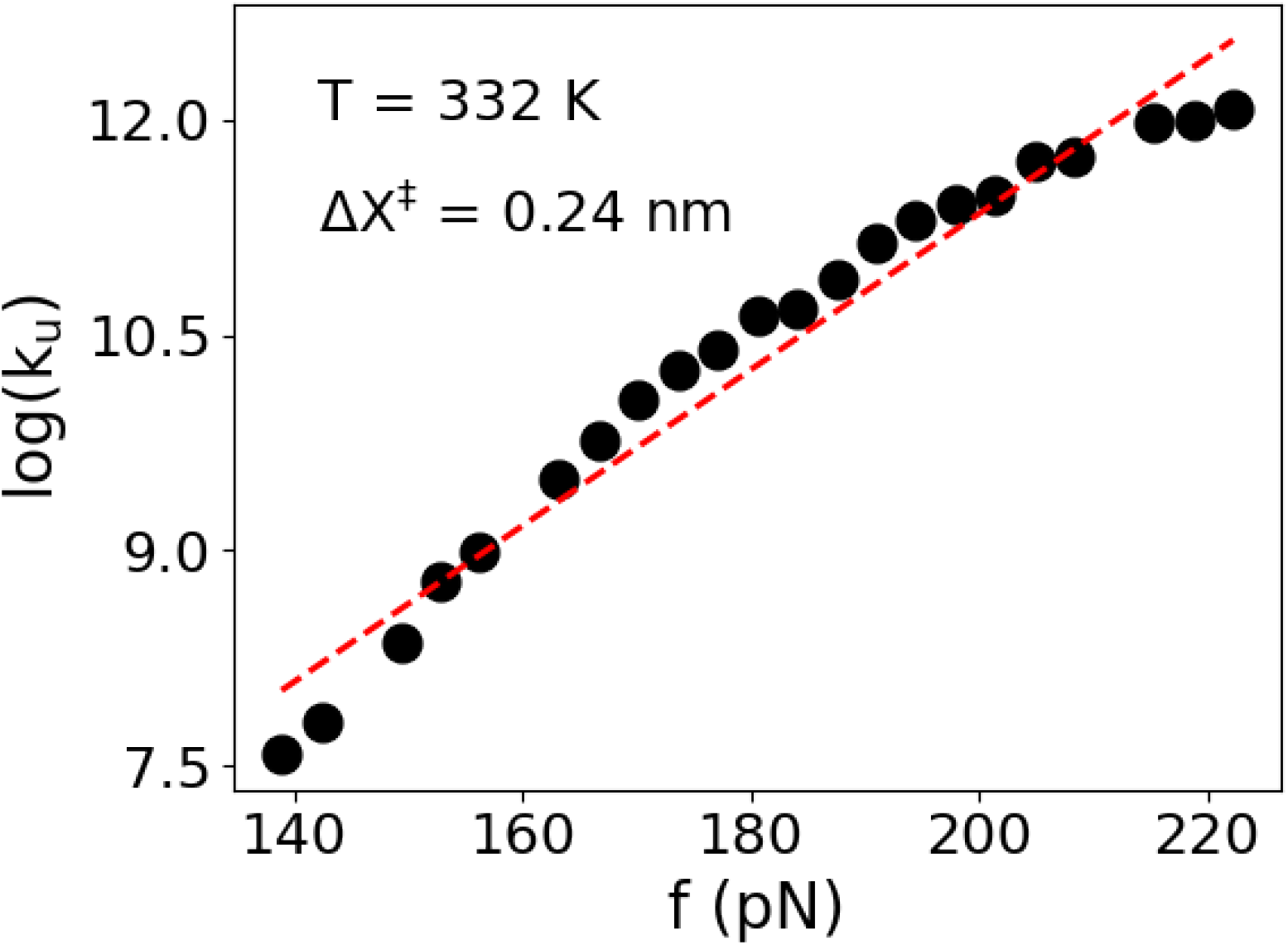
Force dependent unfolding rate, *k_u_*, computed using the force-clamp mode at *T* = 332 K. The dashed line is a fit to the Bell model. The extracted TS location at Δ*X*^‡^ = 0.24 nm is in agreement with the experiments. Deviation from linearity is observed, especially at higher forces leading to a downward curvature.

Although at lower forces (⪅ 170 pN), the plot is linear, deviation from linearity is observed for higher forces leading to a downward curvature. We believe that the downward curvature is due to the weak multidimensionality^26^ (WMD) of the energy landscape (see below). A previous computational study^61^ using an all-atom model also suggested that the Ub unfolding landscape could be multidimensional. In WMD landscapes, *k_u_*(*f*) satisfies the conditions, 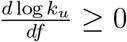 and 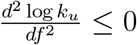, which are the conditions for downward curvature and Ub satisfies them at high *f* values (Figure 1). A minor downward curvature in a WMD energy landscape suggests a force-dependent movement of the TSE (see below).

### Unfolding involves crossing a single major barrier

We also performed force-ramp simulations by pulling one end of Ub with a constant velocity at *T* = 332 K. The average force at which Ub unfolds, ⟨*f_u_*⟩, depends linearly^52,62^ on the logarithm of the loading rate, log(*r_f_*), for loading rates *r_f_* ≤ 8.75 × 10^7^ pN/sec, and deviates from linearity at larger *r_f_* (Figure 2 and S3). Assuming that the Bell model is valid, Evans and Ritchie^63,64^ derived an equation for the most probable unfolding force *f*\*, 
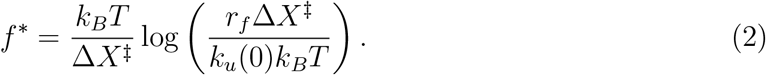

**Figure 2:**
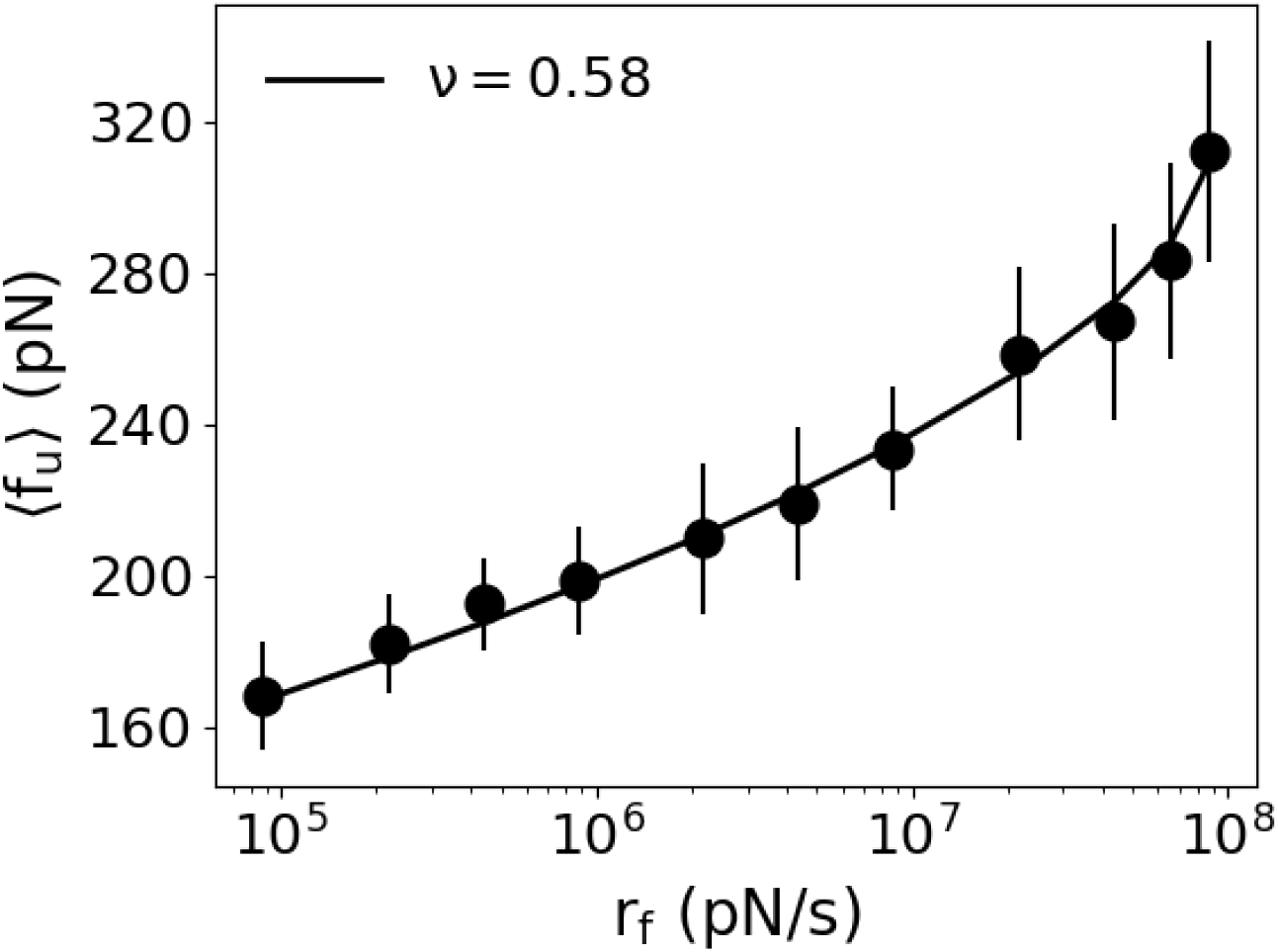
Average unfolding force ⟨*f_u_*⟩ is plotted as a function of log-loading rate log(*r_f_*) at *T* = 332 K. Black solid line is a fit of the data to Eq. 3 and the extracted parameters are *ν* = 0.58±0.0947, Δ*X*^‡^ = 0.66±0.19 nm, Δ*G*^‡^ = 26.07±4.49 *k*_B_*T* and *κ* = 1.265*e* + 07 ± (3.95*e* + 06) s^−1^.

If ⟨*f_u_*⟩ ≈ *f* *, we can estimate Δ*X*^‡^ from the slope of the linear region in the [log *r_f_*, ⟨*f_u_*⟩] plot. The values of Δ*X*^‡^ extracted by fitting only the linear regime in the plots at *T* = 332 K and 300 K are 0.34 nm and 0.38 nm, respectively (Figure S3). These values are in reasonable agreement with Δ*X*^‡^ = 0.24 nm (Figure 1). We also determined the TSE using the *P_fold_* method for the lowest loading rate, *r_f_* = 8.75 × 10^4^ pN/sec, accessible in simulations at *T* = 332 K (Figure S4). The value of Δ*X*^‡^ computed from the TSE is 1.43 nm, and it is in good agreement with the result from constant force simulations (Figure 3), but deviates from the values extracted from the Bell model analysis.

**Figure 3:**
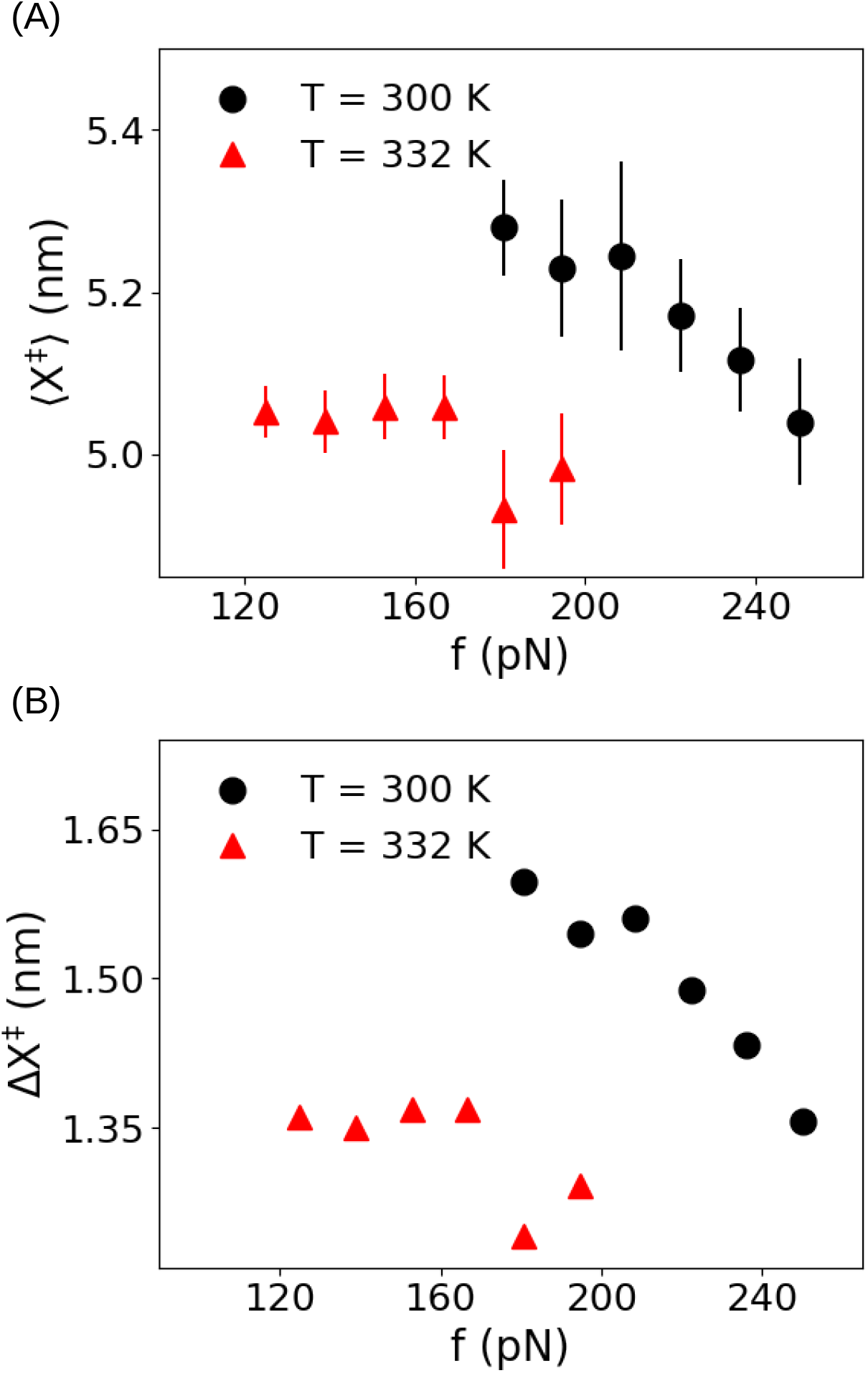
(A) Average end-to-end distance of the Ub conformations in the TSE, ⟨*X*^‡^⟩, as a function of the pulling force *f*. Data shown in circles and triangles are obtained at *T* = 300 K and 332 K, respectively. (B) Relative location of the TS from the native state, Δ*X*^‡^(= ⟨*X*^‡^⟩ − ⟨*X^F^*⟩) as a function of force at *T* = 300 K and 332 K in circles and triangles, respectively. The TS location shifts towards the native state as *f* increases.

The deviation from linearity, with an upward curvature, in the [log *r_f_*, ⟨*f_u_*⟩] plots is evidence for the movement of the TS with force.^65^ To interpret the curvature in log *r_f_* vs. ⟨*f_u_*⟩ plot, a theoretical model^17^ could be used to assess if unfolding occurs by crossing a single barrier or multiple barriers in the one-dimensional free energy profile. In this model, unfolding transitions occur by thermal activation for *f* ≪ *f_c_* and *β*Δ*G*^‡^ ≫ 1, where *f_c_* is the critical force and Δ*G*^‡^ is the free energy of activation. The equation for most probable rupture force is given by,

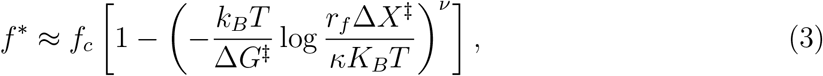
where *κ* is the pre-factor in Kramer’s theory. For unfolding transitions involving a single barrier crossing, with *f* dependent TS, the value of the exponent *ν* is bound by 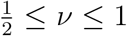, while for transitions with multiple barriers 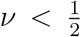. Assuming that ⟨*f_u_*⟩ ≈ *f* *, we fit the data in [log *r_f_*, ⟨*f_u_*⟩] plot using Eq. 3 and obtained *ν* = 0.58 (Figure 2). The relative errors from the fit at each data point are shown in Figure S5. The value *ν* = 0.58 suggests that a single barrier crossing explains the force ramp simulations, which also accords well with the force-clamp data. We conclude that the nonlinearity in the [log *r_f_*, ⟨*f_u_*⟩] plot is due to the force-dependent movement of the TS (see also Figure 3).

### TS location depends on *f*

To check whether the TSE movement follows Hammond postulate, we identified the TSE using the *P_fold_* method^45^ (see Methods section) at *T* = 300 K and 332 K using the constant force simulation data. The average end-to-end distance of the structures in the TSE, ⟨*X*^‡^⟩ varied between ≈ 5.0 - 5.4 nm (Figure 3). The decrease in ⟨*X*^‡^⟩ at the higher temperature is reminiscent of temperature softening demonstrated long ago using AFM experiments on ddFLN4.^66^ For a fixed *T*, the TS shifts to lower ⟨*X*^‡^⟩ values as *f* increases (Figure 3). The average end-to-end distance at *T* = 300 K and 332 K in the absence of force (*f* = 0 pN), calculated from equilibrium simulations are nearly the same ⟨*X^F^*⟩ = 3.68 nm and 3.70 nm, respectively.^49^ The decrease in ⟨Δ*X*^‡^⟩ (= ⟨*X*^‡^⟩ − ⟨*X^F^*⟩) as *f* increases at *T* = 300 K shows that the TSE moves closer to the folded state, which supports Hammond postulate^16,65,67^ (Figure 3). The TSE movement also explains the non-linear behavior in the [*f,* log(*k_u_*(*f*))] plot (Figure 1). We surmise that the gradual TS movement towards the native state with increasing *f* suggests that there is a single major free energy barrier separating the folded and the unfolded states.

The location of the TS (Δ*X*^‡^ ≈ 1.5 nm) at *f* = 195 pN computed directly from the simulations using *P_fold_* analysis, a method that does not rely on a reaction coordinate, differed significantly from the value (Δ*X*^‡^ ≈ 0.24 nm) extracted using Bell fit to either the experimental^41,42^ or the simulation data. The discordance between the Δ*X*^‡^ values indicates that a complete description of the forced-unfolding dynamics requires going beyond one-dimensional free energy profiles.

### *X* alone is not a good reaction coordinate

The disagreement between the Δ*X*^‡^ values extracted using one dimensional free energy profile, with *X* as the reaction coordinate, and computed directly using the *P_fold_* analysis raises the possibility that *X* alone may be an inadequate reaction coordinate for Ub unfolding. If *X* were a good reaction coordinate then the *P_fold_* for the ensemble of structures that are constrained to have *X* ≈ ⟨*X*^‡^⟩ should be ≈ 0.5. At *T* = 332 K and *f* = 125 pN, ⟨*X*^‡^⟩ is 5.1 ± 0.03 nm (Figure 3). We picked 200 Ub conformations whose *X* values are within 5.1 ± 0.03 nm, and performed *P_fold_* analysis for these structures. The distribution, *P* (*P_fold_*), of the *P_fold_* values, shows that a significant fraction of these structures end up in the protein folded state with *P_fold_* = 1. The rest of the structures lead to approximately a uniform distribution between 0 < *P_fold_* < 0.9 (Figure 4). This occurs because most of the Ub conformations with *X* in the range ⟨*X*^‡^⟩ ± *σ* are closer to the folded basin than the TS. As a result, when short pulling simulations using these conformations are initiated, most of these structures reach the folded basin. We conclude that for Ub, *X* alone is not a good reaction coordinate, although both the *f*-dependent unfolding rate and the dependence of the mean unfolding force as a function of *r_f_* do not deviate significantly from the expectation based on an effective 1D energy landscape.

**Figure 4:**
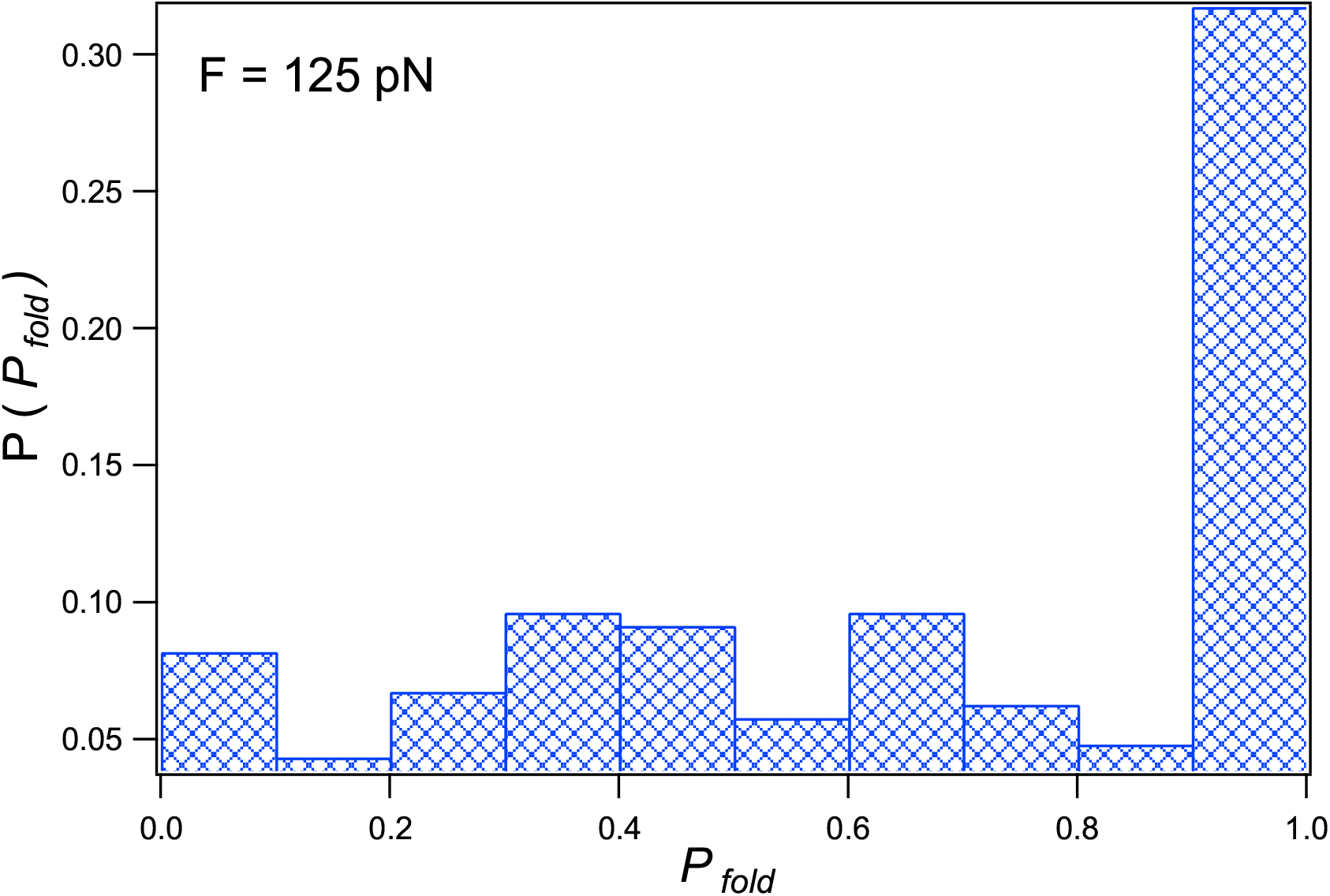
Probability distribution of *P_fold_* values obtained from 200 Ub putative TS structures at *T* = 332 K and *f* = 125 pN. The 200 conformations have *X* within ⟨*X*^‡^⟩ ± *σ* (5.1 ± 0.03 nm). Short pulling trajectories spawned using these structures end up in the folded state basin. The rest of the structures lead to a broad distribution with *P_fold_* between 0 and 0.9. Therefore, from the perspective of the TSE, *X* alone is not an adequate reaction coordinate.

The probability distribution, *P* (*X*), in the absence of *f*, at close to the melting temperature,^49^ *T* = 353 K, where both the folded and unfolded states are sampled, shows that *X* does not resolve the folded and unfolded basins of Ub. There is a cusp separating the two basins in the free energy (Figure 5). This finding is also in agreement with the previous simulation result, ^53^ which reported that *X* could be a poor reaction coordinate for Ub folding. It is possible that a collective coordinate, such as the fraction of native contacts, may be a reasonable choice^49,68^ instead of *X*.

**Figure 5:**
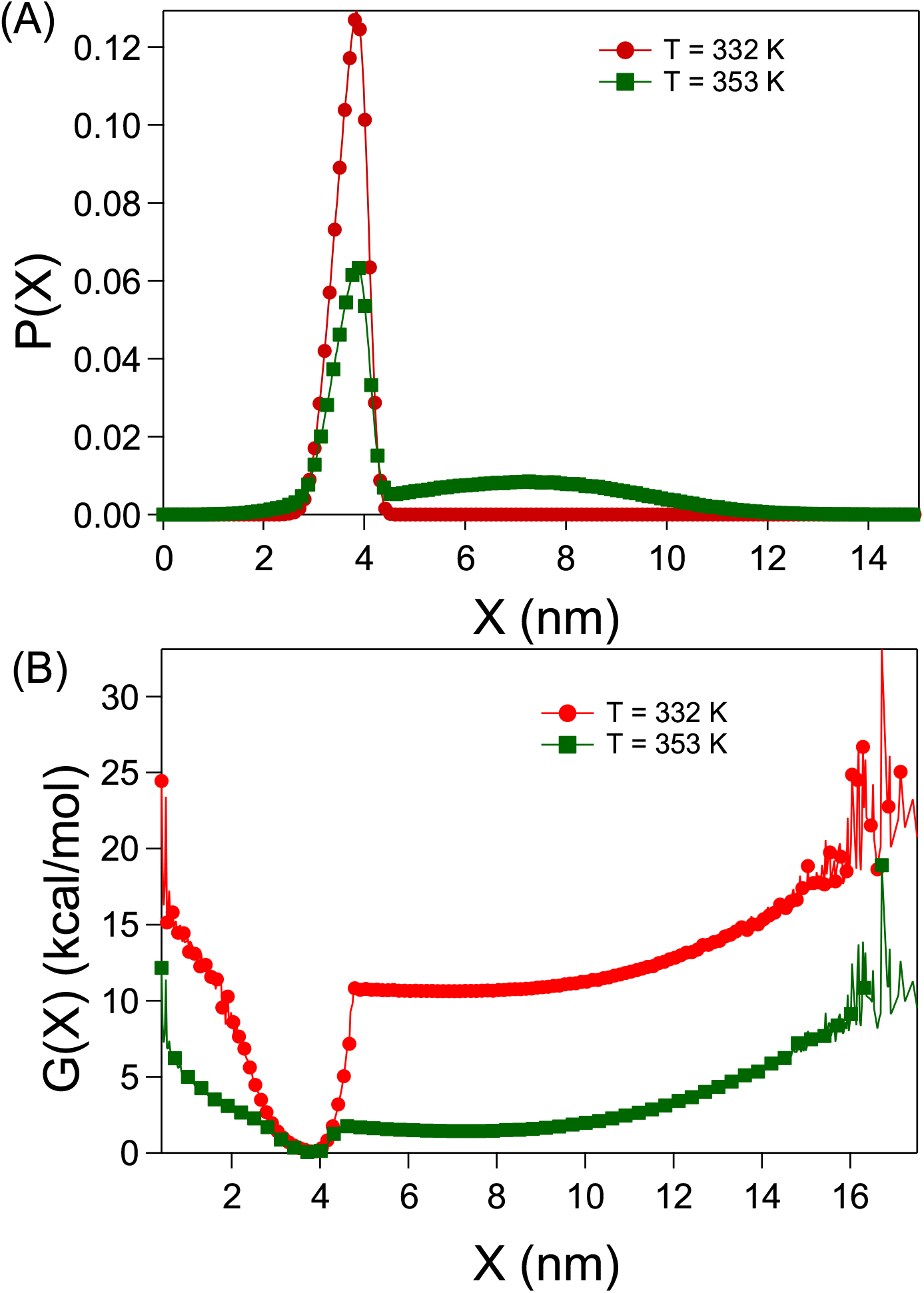
(A) The probability distribution of *X*, *P* (*X*) and (B) FES of Ub projected onto *X*, *G*(*X*) in the absence of external force (*f* = 0 pN) at *T* = 332 K and 353 K. The melting temperature of Ub is ≈ 353 K.

## Conclusions

A major advantage of single-molecule pulling experiments is that the energy landscape of proteins can be measured without altering the intra-protein or protein solvent interactions. However, we need useful theoretical models in order to extract the relevant kinetic properties related to the protein’s structural transitions from the experimental data. The common assumptions in the most fruitful models are: (1) Protein unfolding can be described as a barrier crossing in an effective one dimensional energy landscape, (2) The location of the transition state is not very sensitive to the applied force. These assumptions need to be tested to ensure that extracted quantities are physically meaningful, and consistent with independent experiments and/or reliable simulations.

Unfolding Ub by force, which has been extensively studied by experiments, is a curious case. The dependence of the unfolding rates on force and the mean rupture force as a function of loading rates only exhibit minor deviations from theoretical predictions based on 1D energy landscape. On the other hand, the structures that are generated by constraining the extension to the TS location extracted by fitting the simulation data to the one-dimensional analytical models do not satisfy the *P_fold_* ≈ 0.5 requirement. By combining both these findings, we demonstrate that the energy landscape of Ub is weakly multidimensional in which the location of the transition state moves in response to the applied force. Our study shows that only by combining analysis of experimental data using simple theories together with reliable simulations can one learn about the unfolding energy landscape, especially for systems which exhibit weak multidimensionality.

## Acknowledgement

A part of this work is funded by the grant to G.R. by the National Supercomputing Mission (MeitY/R&D/HPC/2(1)/2014). D.T. acknowledges grants from the NIH (GM - 107703), and the Welch Foundation (Grant F-0019) through the Collie-Welch chair. The computations are performed using the TUE and Cray XC40 clusters at IISc.

## Supporting Information Available

Description of the coarse-grained self-organized polymer-side chain protein model and simulation method; Tables S1-S3; Figures S1-S5. Table-S1: SOP-SC energy parameters. Table-S2: SOP-SC model parameters for ubiquitin. Table-S3: Side-chain radii of amino acids. Figure-S1: Multiple independent trajectories showing two-state unfolding of Ub. Figure-S2: Ub dominant unfolding pathway. Figure-S3: Average unfolding force ⟨*f_u_*⟩ as a function of the loading rate ln(*r_f_*). Figure-S4: *P_fold_* analysis to identify the TSE from force ramp simulations. Figure-S5: Relative errors from the fit of Eq. 3 to the [log(*r_f_*), ⟨*f_u_*⟩] data.

