## Supporting Information (SI) for "Energy Landscape of Ubiquitin is Weakly Multidimensional"

### Methods

#### Self Organized Polymer-Side Chain (SOP-SC) model:

We have used the SOP-SC model[1,2] to perform pulling simulations on Ubiquitin (Ub). We showed previously that the simulations using the SOP-SC model are remarkably accurate in capturing the temperature and denaturant effects on Ub folding.[3,4] In the SOP-SC model, each amino acid is represented by two beads. The backbone atoms of an amino acid are represented by a bead positioned at the center of $C_\alpha$ atom, and the side chain atoms are represented by another bead positioned at the center of mass of the side-chain. The SOP-SC model for Ub is constructed using the protein data bank (PDB) structure with PDB ID: 1UBQ.[5] Missing hydrogen atoms are added to the structures using the program visual molecular dynamics (VMD)[6] before calculating the center of mass of the side-chain. The SOP-SC energy function is described in terms of bonded ($E_B$) and non-bonded interactions ($E_{NB}$). Covalently connected beads interact via a bonded potential ($E_B$). Non-bonded interactions ($E_{NB}$) consist of native (N) and non-native (NN) interactions. Interactions between two beads are considered native, if they are separated by at least three bonds and are within a cut-off distance ($R_c$) in the SOP-SC model of the PDB structure. Any other non-covalent interactions are considered as non-native interactions ($E_{NN}$). Native interactions between neighboring side-chain beads are ignored because of their close proximity in both folded and unfolded states. The SOP-SC energy function for a protein conformation described by the set of coordinates $\{\mathbf{r}\}$ is given by,

$$E_{CG}(\{\mathbf{r}\}) = E_B + E_{NB}^{N} + E_{NB}^{NN}. \tag{S1}$$

The bonds between two consecutive beads are modeled using the finite extensible nonlinear elastic (FENE) potential by,

$$E_B = -\sum_{i=1}^{N_B} \frac{k}{2} R_0^2 \log\Big(1 - \frac{(r_i - r_{cry,i})^2}{R_0^2}\Big), \tag{S2}$$

where $N_B$ is the total number of bonds present between the covalently linked beads, $r_i$ is the distance between $i^{th}$ pair of beads and $r_{cry,i}$ is the distance between the same $i^{th}$ pair of beads in the SOP-SC representation of the PDB structure. The native interactions $E_{NB}^{N}$, are modeled using a Lennard-Jones type of potential given as,

$$\begin{aligned} E_{NB}^{N} = & \sum_{i=1}^{N_N^{bb}} \epsilon_h^{bb} \big[ \big(\frac{r_{cry,i}}{r_i}\big)^{12} - 2\big(\frac{r_{cry,i}}{r_i}\big)^{6} \big] + \sum_{i=1}^{N_N^{bs}} \epsilon_h^{bs} \big[ \big(\frac{r_{cry,i}}{r_i}\big)^{12} - 2\big(\frac{r_{cry,i}}{r_i}\big)^{6} \big] \\ & + \sum_{i=1}^{N_N^{ss}} 0.5 \times 300 k_B \times (0.7 - \epsilon_i^{ss}) \big[ \big(\frac{r_{cry,i}}{r_i}\big)^{12} - 2\big(\frac{r_{cry,i}}{r_i}\big)^{6} \big], \end{aligned} \tag{S3}$$

where $N_N^{bb}$, $N_N^{bs}$ and $N_N^{ss}$ represent the total number of native contact pairs present between backbone-backbone, backbone-side chain, and side chain-side chain beads, respectively. $k_B$ is the Boltzmann constant, $r_i$ is the distance between $i^{th}$ pair of beads and $r_{cry,i}$ is the distance between the same $i^{th}$ pair of beads in the SOP-SC representation of the PDB structure. $\epsilon_h^{bb}$ and $\epsilon_h^{bs}$ denote the strength of interaction between backbone beads and backbone-side chain beads respectively. Strength of interactions between the side-chain beads $\epsilon_i^{ss}$ are obtained from the Betancourt-Thirumalai statistical potential.[7]

Non-native interactions ($E_{NB}^{NN}$) are modeled as,

$$E_{NB}^{NN} = \sum_{i=1}^{N_{NN}} \epsilon_l \big(\frac{\sigma_i}{r_i}\big)^6 + \sum_{i=1}^{N_{ang}^{bb}} \epsilon_l \big(\frac{\sigma^{bb}}{r_i}\big)^6 + \sum_{i=1}^{N_{ang}^{bs}} \epsilon_l \big(\frac{\sigma_i^{bs}}{r_i}\big)^6, \tag{S4}$$

where $N_{NN}$ is the total number of non-native interaction pairs present in the SOP-SC model, $\sigma_i$ is sum of the radii of the beads in $i^{th}$ pair of non-native interactions. $\sigma^{bb}$ is the diameter of the backbone beads, and $\sigma_i^{bs} (= f[\sigma^{bb} + \sigma_i^{sc}]/2.0)$ is the sum of the radii of the backbone and the side chain in the $i^{th}$ pair of angular interactions scaled by a factor $f = 0.8$. The second and third terms in eq. S4 model the bond angle potential between beads separated by two bonds. $N_{ang}^{bb}$ and $N_{ang}^{bs}$ are the total number of bond angles between the backbone beads and between the backbone-side chain beads. The values of the parameters in the SOP-SC energy functions and the side-chain radii are listed in Table S1, S2 and S3, respectively.

#### Simulations:

To provide a realistic description of Ub unfolding kinetics in the presence of external force, $f$, we performed Brownian dynamics simulations. The equations of motion were integrated using the Ermak-McCammon algorithm,[8] $\vec{r_i}(t+h) = \vec{r_i}(t) + \frac{h}{\zeta}\vec{F_c} + \vec{\Gamma}$, where $\vec{r_i}(t)$ is the position of the bead $i$ at time $t$, $\zeta$ is the friction coefficient, $\vec{F_c} = \frac{\partial E_{TOT}}{\partial \vec{r_i}}$, $\vec{\Gamma}$ is the random force with a Gaussian distribution with mean zero and variance $\langle \Gamma(h)^2 \rangle = \frac{2k_BTh}{\zeta}$, and $T$ is the temperature. The friction coefficient $\zeta = 50\ m/\tau_H$ approximately corresponds to the value in water and $h = 0.005\ \tau_H$ is the time step used to advance the simulation. In the simulations, the characteristic unit of length $a = 1$ Å, energy $\epsilon = 1$ kcal/mol, and mass $m = 1.8 \times 10^{-22}$ g (typical mass of the bead). The unit of time in the simulations is $\tau_L(= \sqrt{ma^2/\epsilon}) = 0.51$ ps. In Brownian dynamics, simulation time is mapped into real time, $\tau_H$ using $\tau_H \approx \frac{\zeta_H a^2}{k_BT} = \frac{(\zeta_H \tau_L/m)\epsilon}{k_BT}\tau_L \approx 43$ ps.

In the force-clamp simulations where unfolding is initiated by applying a constant force, $f$, we fixed the position of the $N$-terminal end of the protein (backbone bead of residue 1), and applied a constant $f$ on the $C$-terminal end (backbone bead of residue 76). In the force-ramp simulations, the backbone bead of residue-1 is fixed, and the backbone bead of residue-76 is attached to a spring, with a spring constant, $k_s$= 35 pN/nm. The other end of the spring is moved with a constant velocity, $v$, to ramp up the force to unfold the protein. We used different values of the velocities leading to loading rates, $r_f = k_s v$, in the range 8.75 $\times 10^4$ pN/s to 8.75 $\times 10^7$ pN/s to unfold the protein.

**Fitting Procedure:** We used maximum-likelihood estimation (MLE) to determine the optimum fitting parameters, relative position of the transition state ($\Delta X^{\ddagger}$) and intrinsic unfolding rate ($k_u(0)$), from the force-clamp data using the Bell model (Eq. 1). For $M$ constant forces $f_i$ with $N_i$ measurements of unfolding time, average unfolding time $\tau_i$, and force dependent unfolding rate $k(f_i)$, the log likelihood function is defined as $L = \sum_{i=1}^{M} N_i(\log k(f_i) - k(f_i)\tau_i)$.[9] The maximum value of $L$ is found by minimizing the negative value of $L$ using Limited-memory Broyden-Fletcher-Goldfarb-Shanno with bound constraints

(L-BFGS-B) algorithm on the data set $\{f_i, N_i, \tau_i\}$. The extracted parameters are used to fit the $[\log k_u(f), f]$. For the force-ramp data, average unfolding force $\langle f_u \rangle$ is fitted as a function of log-loading rate $\log(r_f)$ following Eq. 3 using non-linear least-square method. We used SciPy (version 1.6.0) for maximizing the log likelihood function and implementing the non-linear least-square method. Errors in fitting parameters are computed from the respective covariance matrices.

Table S1: SOP-SC energy parameters

| Parameters | Protein |
|---|---|
| $R_o$ | 2.0 Å |
| $k$ | 20 kcal/mol/Å$^2$ |
| $R_c$ | 8 Å |
| $\epsilon_h^{bb}$ | 0.5 kcal/mol[a] |
| $\epsilon_h^{bs}$ | 0.5 kcal/mol[a] |
| $\epsilon_l$ | 1.0 kcal/mol[a] |
| $\sigma^{bb}$ | 3.8 Å |
| $\epsilon$ | 10.0 |

[a] Values are chosen such that the protein melting temperature in simulations is approximately in agreement with the experiments.[10,11]

Table S2: SOP-SC model parameters for ubiquitin

| Parameters | 1UBQ |
|---|---|
| $N_B$ | 151 |
| $N_N^{bb}$ | 177 |
| $N_N^{bs}$ | 486 |
| $N_N^{ss}$ | 204 |
| $N_{NN}$ | 10159 |
| $N_{ang}^{bb}$ | 74 |
| $N_{ang}^{bs}$ | 150 |

Table S3: Side-chain radii of amino acids

| Residue | Radius (Å) |
|---|---|
| Gly | 0.5 |
| Ala | 2.52 |
| Val | 2.93 |
| Leu | 3.09 |
| Ile | 3.09 |
| Met | 3.09 |
| Phe | 3.18 |
| Pro | 2.78 |
| Ser | 2.59 |
| Thr | 2.81 |
| Asn | 2.84 |
| Gln | 3.01 |
| Tyr | 3.23 |
| Trp | 3.39 |
| Asp | 2.79 |
| Glu | 2.96 |
| Hsd | 3.04 |
| Lys | 3.18 |
| Arg | 3.28 |
| Cys | 2.74 |

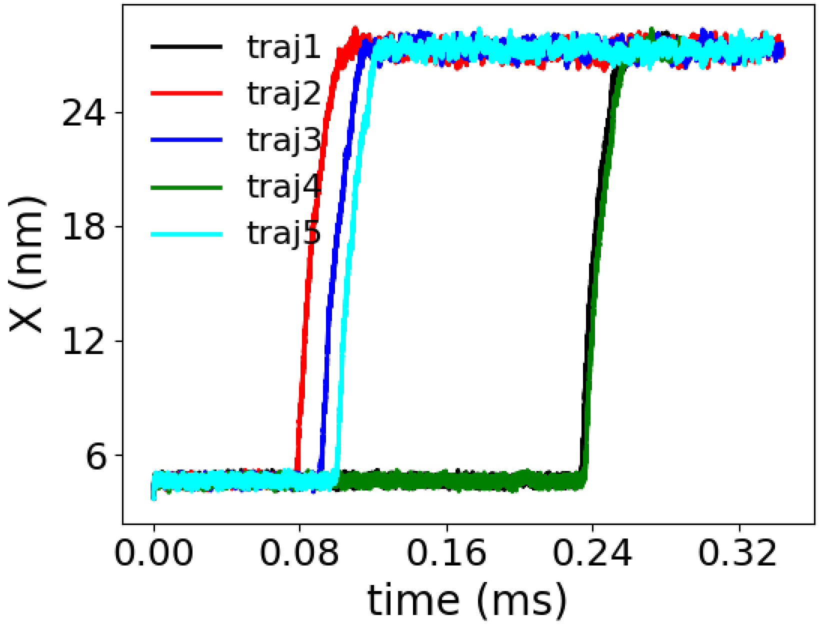


Figure S1: Two-state unfolding transition is observed in Ub. Five independent constant force pulling simulations are shown in different colors for $f = 149$ pN and $T = 332$ K.

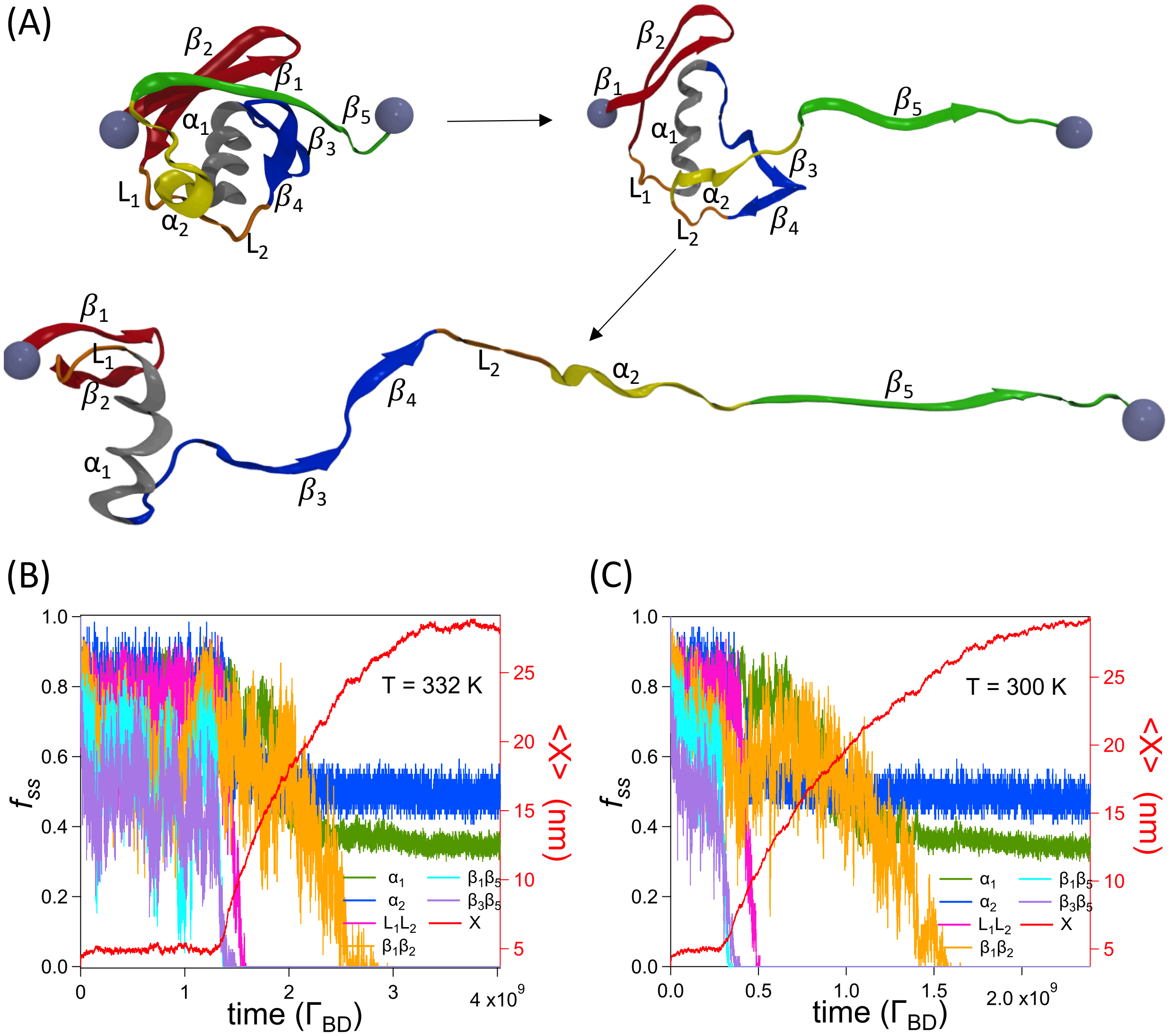


Figure S2: Dominant Ub unfolding pathway in the presence of external force at $T = 332$ K. (A) Strand $\beta_5$ detaches first disrupting its contacts with $\beta_1$ and $\beta_3$. Following this, the contacts between the loops $L_1L_2$ fall apart destabilizing the remaining structure. In the last stages, the $N$-terminal hairpin $\beta_1\beta_2$ unfolds. The same unfolding pathway is observed at $T = 300$ K. Fraction of native contacts in various secondary structural elements $f_{ss}$ and $X$ as a function of time at (B) $T = 332$ K; $f = 208$ pN and (C) $T = 300$ K; $f = 222$ pN. $X$ as a function of time is shown in red, and the scale is on the right.

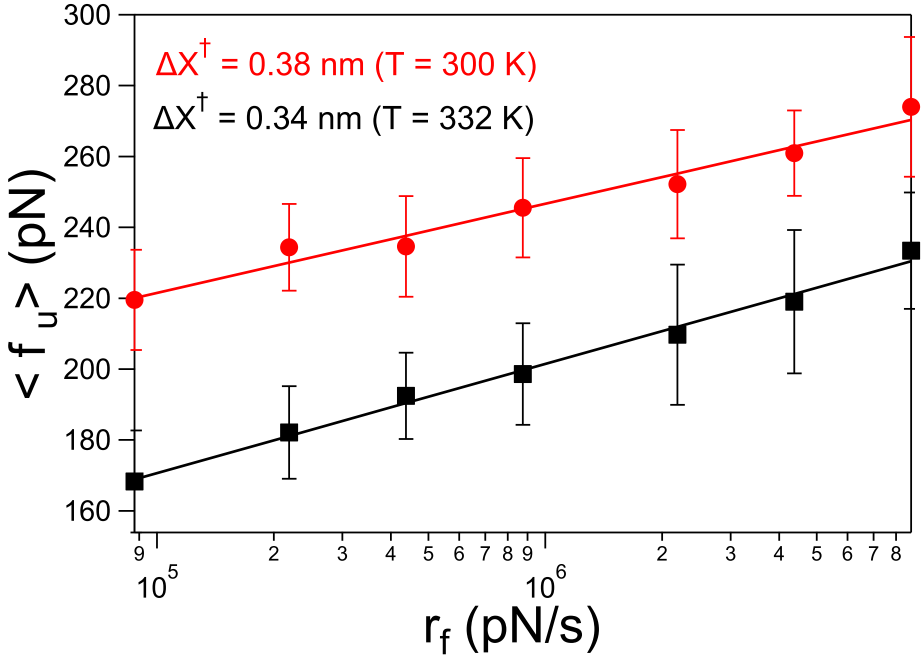


Figure S3: Average unfolding force $\langle f_u \rangle$ depends linearly on the logarithm of the loading rate $\ln(r_f)$ for lower loading rates in the force-ramp experiments. The simulations are performed at $T = 332$ K. The line is a linear fit to extract the slope $k_B T/\Delta X^{\ddagger}$.

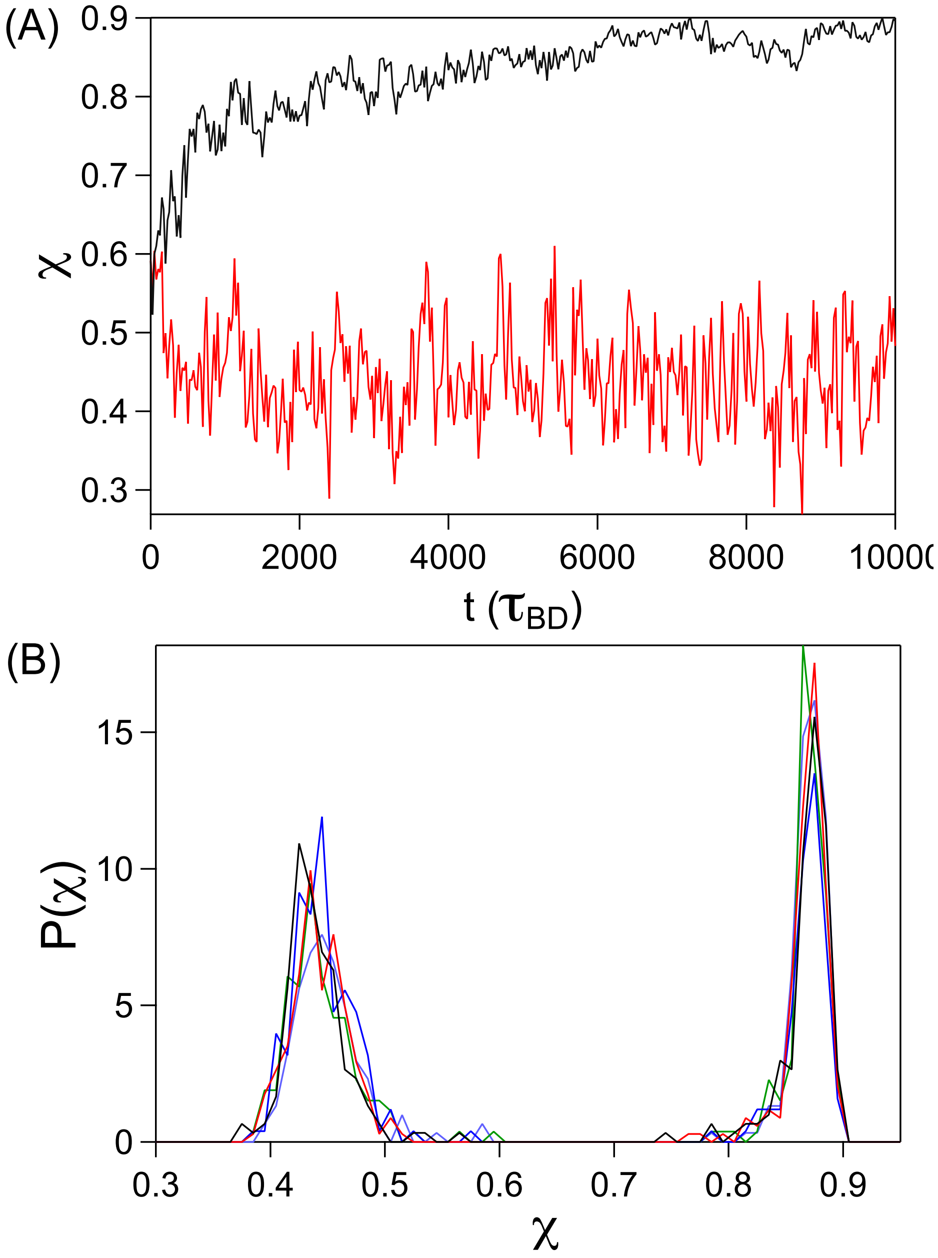


Figure S4: (A) The structural overlap function $\chi$ is plotted as a function of time $t$ for trajectories spawned from the Ub structures in the TSE. The trajectories end up in both the folded basin (red) and the unfolded basin (black). (B) The distribution of the final structural overlap parameter, $\chi$, calculated from 500 simulation trajectories spawned using the TSSs as the initial starting structure. Data are shown for five different structures. The distribution shows that roughly half of these trajectories go into the folded basin and the other half into the unfolded basin, demonstrating that these protein conformations are the TSSs. The data is from force-ramp simulations performed at $T = 332$ K with a loading rate, $r_f = 8.75 \times 10^4$ pN/sec.

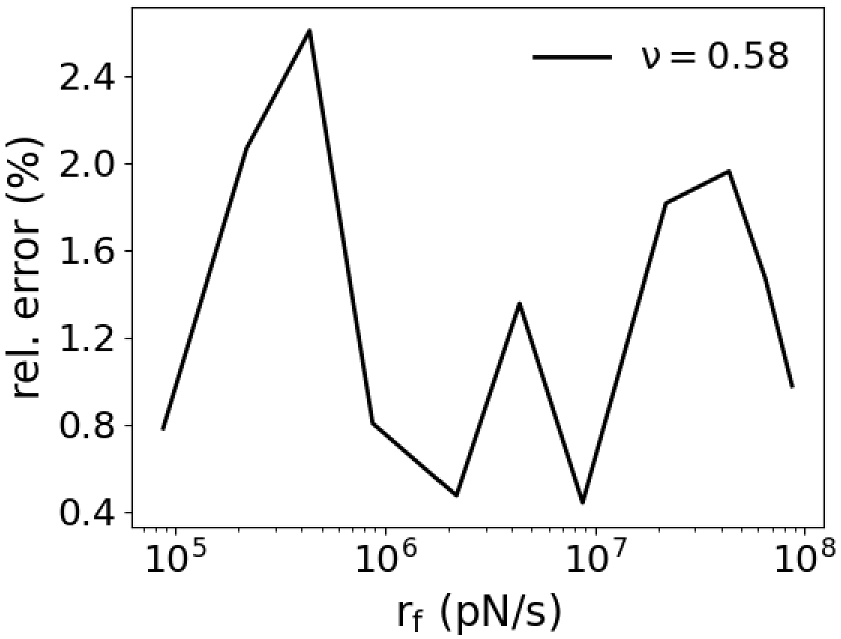


Figure S5: Relative error from the fit at each data point is computed using $(|f^*_{fit} - f^*_{exp}|/f^*_{exp} \times 100)$, where $f^*_{fit}$ and $f^*_{exp}$ are values from the fit and simulation, respectively. The quality of the fit is given by $\chi^2_{red} = 0.08$ and $\chi^2 = 0.56$, where $\chi^2 = \sum_i \frac{(O_i - E_i)^2}{E_i}$ and $\chi^2_{red} = \frac{\chi^2}{n-m}$. Here $O_i$ is the observed value and $E_i$ is the expected value of the $i^{th}$ data from the fit, $n$ is the total number of observations and $m$ is the total number of fitted parameters
